# Magnesium as a Conformational Gatekeeper of KRAS: Structural Dynamics and Therapeutic Implications

**DOI:** 10.64898/2026.02.20.707046

**Authors:** Bindu Y. Srinivasu, Tanvi S. Damerla, Alexander Stec, Zhiwei Zhou, John R. Engen, Kenneth D. Westover, Thomas E. Wales

## Abstract

Magnesium serves as an essential cofactor for small GTPases, yet its structural role in regulating KRAS conformational dynamics and nucleotide exchange remains poorly understood. Here, we combine hydrogen–deuterium exchange mass spectrometry (HDX-MS), native mass spectrometry, and functional assays to elucidate how Mg^2+^ stabilizes the KRAS conformational ensemble and constrains transitions between GDP- and GTP-bound states. Depletion of Mg^2+^ triggers widespread increases in structural dynamics throughout KRAS—spanning the p-loop, α1-helix, switch I, nucleotide-binding region, and distal helices—revealing a global loosening of the protein fold that favors an open, nucleotide exchange-competent state. Mg^2+^ titration experiments demonstrate that individual structural elements exhibit distinct Mg^2+^ dependencies: the p-loop and α1-helix recover native dynamics at micromolar concentrations, whereas switch I requires millimolar levels, underscoring its exceptionally high sensitivity to Mg^2+^ for structural stabilization. KRAS bound to the catalytic domain of exchange factor SOS1 displays an HDX signature closely resembling the Mg^2+^-free state, indicating that SOS1 promotes nucleotide exchange by transiently perturbing Mg^2+^ coordination while simultaneously stabilizing switch I. Consistently, phosphomimetic KRAS S17E variant, which disrupts a critical Mg^2+^-coordinating residue, exhibits pronounced global destabilization—reinforcing the central importance of Mg^2+^ in maintaining structural integrity. Taken together our findings show that Mg^2+^ acts as a master regulator of KRAS structural dynamics and reveal Mg^2+^-sensitive hotspots that might represent promising targets for next-generation KRAS therapeutics.

## Introduction

Divalent metal ions—including Fe^2+^, Ca^2+^, Zn^2+^, and Mg^2+^—serve as ubiquitous regulators of protein structure and function throughout biology. These cations participate in enzymatic catalysis, coordinate ligand binding, provide electrostatic stabilization, and control conformational states across diverse protein families [1–3]. Among them, magnesium occupies a uniquely central position: it is the predominant cofactor required for nucleotide binding, phosphoryl-transfer reactions, and maintaining conformational fidelity in numerous GTP- and ATP-dependent proteins [3, 4]. Through coordinating phosphate groups, modulating local electrostatics, and influencing conformational flexibility, Mg^2+^ underpins fundamental cellular processes including ATP-dependent reactions, RNA metabolism, and intracellular signaling [5]. While total cellular Mg^2+^ concentrations range from 15–18 mM, the free Mg^2+^ pool available for biochemical reactions is approximately 0.8–1.2 mM [6]. Despite this abundance, the precise structural mechanisms by which Mg^2+^ shapes protein conformational landscapes and particularly its role in regulatory switching remain incompletely defined.

The RAS family of small GTPases exemplifies this knowledge gap. As central signaling regulators and among the most consequential oncogenes in human cancer [7, 8], RAS proteins critically depend on Mg^2+^ for proper function. The RAS family comprises four closely related genes—NRAS, HRAS, KRAS4A, and KRAS4B, together harbor activating mutations in over 25% of human cancers. KRAS predominates among these, accounting for approximately 86% of all RAS alterations [9]. Functioning as a molecular switch, KRAS cycles between an inactive GDP-bound state and an active GTP-bound state [10]. This nucleotide-dependent switching mechanism enables KRAS to orchestrate key signal transduction pathways—including the Ras–Raf–MEK–ERK and Ras–PI3K–Akt cascades—that govern cellular growth, proliferation, and survival [11–13]. The guanine nucleotide exchange factor (GEF), SOS1 catalyzes KRAS activation through nucleotide exchange; specifically, its catalytic CDC25 domain accelerates GDP release and enables GTP loading [14]. Conversely, GTPase-activating proteins (GAPs) stimulate the intrinsic GTP hydrolysis of KRAS, returning it to the inactive state and terminating downstream signaling [15, 16].

The clinical significance of KRAS mutations—present in 20–30% of all cancers, with particularly high frequencies in lung, colorectal, and pancreatic tumors [17, 18] has spurred intensive efforts to elucidate both structural and regulatory mechanisms underlying KRAS function, with the ultimate goal of enabling structure-based drug design. Identifying exploitable vulnerabilities in these mechanisms could yield therapeutics effective across diverse oncogenic KRAS mutants. Structural analyses reveal that all RAS proteins share a highly conserved G-domain (residues 1–166) alongside a more divergent hypervariable region at the C-terminus (HVR; residues 167–188/189) that facilitates membrane localization through post-translational modifications. The N-terminal effector lobe (residues 1–86) containing the key regulatory elements—the p-loop (residues 10–17), switch I (residues 25–40), and switch II (residues 57–75)—and a allosteric lobe (residues 87–166) harboring the nucleotide-binding region (residues 114–122) [19, 20]. Critically, KRAS requires a divalent Mg^2+^ cation as a cofactor for proper function (Figure 1B) [21]. Mg^2+^ coordinates directly with Ser17, the β-phosphate of GDP (or with Thr35 and both β- and γ-phosphates when GTP is bound), with the remaining octahedral coordination sphere occupied by water molecules and contributions from switch I residues Tyr32, Asp33, Pro34, and Ile36 [22].

**Figure 1:**
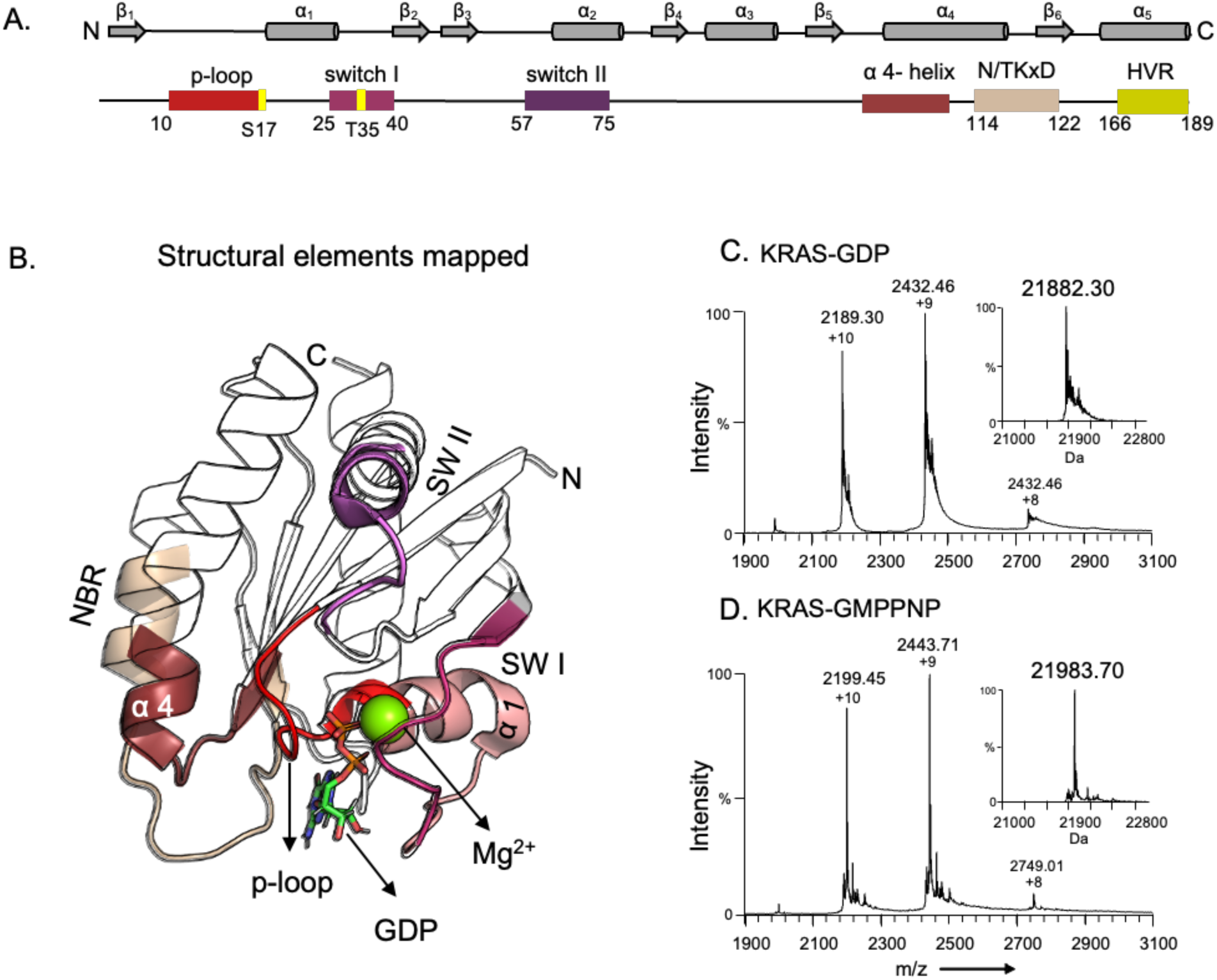
Structural elements of KRAS. (A) Secondary structure diagram of the KRAS protein, highlighting key functional regions: the p-loop, switch I, switch II, α4-helix, N/TKXD motif, and the hypervariable region (HVR), along with specific residue numbers. (B) Cartoon representation of the KRAS protein (PDB ID: 4OBE) bound to GDP and Mg^2+^. The structural elements highlighted in panel A are mapped onto this 3D structure. Native mass spectrometry of KRAS in GDP and GMPPNP-bound states. (C, D) ESI mass spectra of KRAS where the x-axis shows the m/z ratio and the y-axis represents the intensity. Each spectra shows three different charge states (+8, +9 and +10 charge states) and the inset shows the deconvoluted mass obtained from the transformation of the charge states for each spectrum using MaxEnt1from MassLynx 4.1 (Waters). (C) KRAS bound to GDP with mass of 21882.3 Da (KRAS alone = 21,439.1 Da and 443.2 Da for GDP). (D) KRAS-bound to GMPPNP with a mass of 21,983.7 Da (KRAS alone = 21,439.1 Da and 544.6 Da for GMPPNP)

Prior NMR studies by Pálfy *et al*. demonstrated that KRAS samples a millisecond-timescale two-state exchange in both GDP- and GTP-loaded forms, with a sparsely populated “open conformation” in the GDP state resembling a nucleotide-bound, Mg^2+^-free structure characterized by shortened β2–β3 strands and partial release of switch I [23]. Consistent with this, X-ray crystallographic studies revealed that KRAS constructs lacking the initiator methionine and N-terminal acetylation adopt Mg^2+^-free conformations featuring a flexible N-terminus that destabilizes the interswitch region and drives switch I into an open, exchange-ready state [22]. These observations parallel structural findings from KRAS–SOS1 complexes captured by cryo-EM, wherein KRAS appears devoid of Mg^2+^, nucleotide-free conformation [24]. Molecular dynamics simulations have further illuminated the mechanism, revealing that SOS1 catalyzes nucleotide release by displacing switch I and switch II from the nucleotide-binding pocket while repositioning Ala59 into the site normally occupied by Mg^2+^, thereby disrupting metal coordination and destabilizing GDP binding [14]. Yet despite these structural snapshots from crystallography, cryo-EM, and protein dynamics through computational simulations, how Mg^2+^ shapes the global and local structural dynamics of KRAS—particularly given the protein’s inherently dynamic nature—has remained largely unexplored.

Hydrogen–deuterium exchange mass spectrometry (HDX-MS) offers a powerful approach for analyzing protein structural dynamics by monitoring the substitution of backbone amide hydrogens with deuterium [25–28]. Both the extent and rate of deuterium incorporation report on multiple structural features, including backbone solvent accessibility, hydrogen-bond strength, and conformational flexibility. Using HDX-MS, we demonstrate here that Mg^2+^ depletion disrupts not only the immediate coordination pocket but also allosterically propagates increased backbone dynamics to distal regulatory elements—establishing Mg^2+^ as a global stabilizer rather than merely a local cofactor. Furthermore, our data reveal that while the SOS1 catalytic domain perturbs Mg^2+^ coordination to promote nucleotide exchange, it simultaneously preserves key structural features, functioning as both a catalytic activator and a molecular “chaperone”. Together, these findings advance beyond earlier static and fragmentary views of KRAS structural behavior, providing a comprehensive solution-phase map of how Mg^2+^ governs the conformational landscape and activation mechanism of KRAS.

## Results

### Mg^2+^ Depletion Facilitates Nucleotide Exchange

To assess the structural consequences of Mg^2+^ depletion for KRAS nucleotide exchange, first we employed native mass spectrometry (native-MS) to monitor nucleotide occupancy through alterations in molecular mass. In native-MS, proteins in aqueous ammonium acetate enter the mass spectrometer operated under optimized conditions designed to preserve noncovalent interactions and in near-native conditions, enabling direct detection of intact protein complexes [29]. We first incubated KRAS–GDP with excess EDTA to chelate and to deplete Mg^2+^, then added an excess of the non-hydrolyzable GTP analog GMPPNP to initiate exchange. Following the addition of excess MgCl₂ to sequester EDTA, samples were exchanged into buffer containing 50 mM ammonium acetate (pH 7.8) for analysis. Native-MS spectra revealed that untreated KRAS–GDP exhibited a molecular mass of 21,882.3 ± 0.49 Da (Figure 1C; corresponding to 21,439.1 Da for KRAS plus 443.2 Da for GDP), whereas the EDTA-treated, GMPPNP-loaded sample displayed a mass of 21,983.7 ± 0.21 Da (21,439.1 Da for KRAS plus 544.6 Da for GMPPNP; Figure 1D). This unambiguous mass shift provides direct evidence corroborating earlier biochemical findings that Mg^2+^ depletion facilitates nucleotide exchange [30], demonstrating that Mg^2+^ depletion promotes GDP dissociation.

### Mg^2+^ Clamps the Nucleotide-Bound Conformation of KRAS

To investigate how Mg^2+^ coordination contributes to KRAS structural stability and conformational dynamics, we performed HDX-MS on recombinant KRAS protein in both Mg^2+^-bound and EDTA-treated Mg^2+^-deplete states. In HDX-MS, protein is diluted into a D₂O-based buffer and labile hydrogens (both in the backbone and side chains) undergo exchange with solvent deuterium; only backbone amide hydrogens are monitored under LC MS experimental conditions. Those amide hydrogens that are buried, hydrogen-bonded, or otherwise structurally protected exchange slowly, whereas regions rendered solvent-accessible through intrinsic motions or allosteric rearrangements exchange more rapidly. Deuterium uptake patterns thus provide a sensitive, quantitative readout of local stability, dynamic behavior, and conformational changes throughout the protein. Here, we generated Mg^2+^-deplete KRAS by treating purified protein with a 50-fold molar excess of EDTA to chelate bound Mg^2+^, followed by buffer exchange into 20 mM HEPES, 150 mM NaCl (pH 7.8) to remove residual chelator.

KRAS samples in GDP-bound (inactive) and GMPPNP-bound (active) forms, prepared in both Mg^2+^-replete and Mg^2+^-depleted states, were exposed to 18-fold excess of D₂O labeling buffer (20 mM HEPES, 150 mM NaCl, pD 7.8). At defined time points, the deuterium exchange reaction was quenched, protein was digested online using Nepenthesin II, and deuterium incorporation was quantified for each peptide by ESI-MS. In Mg^2+^-bound KRAS–GDP, the p-loop (residues 10–19), α1-helix (residues 17–30), α4-helix (residues 80–90), nucleotide-binding region (residues 114–132), and C-terminal region (residues 137–159) exhibited low initial deuterium uptake (10–30%) with modest increases at later time points, reflecting stable, solvent-protected conformations. In contrast, switch I (residues 31–40) and switch II (residues 57–75) displayed markedly higher exchange rates. Switch I showed rapid deuterium incorporation, beginning at approximately 30% at the earliest time point and exceeding 50% by 10 minutes, whereas switch II reached approximately 30% over the same period (Figure 2A) consistent with the established dynamic character of these regulatory switch regions [31]. Mg^2+^-bound KRAS–GMPPNP displayed broadly similar patterns across most regions, though switch II and the nucleotide-binding region exhibited substantially higher initial deuterium incorporation (exceeding 50% initially and rising above 60% at later time points; Figure 2C), likely reflecting the enhanced conformational mobility associated with the GTP-like active state.

**Figure 2:**
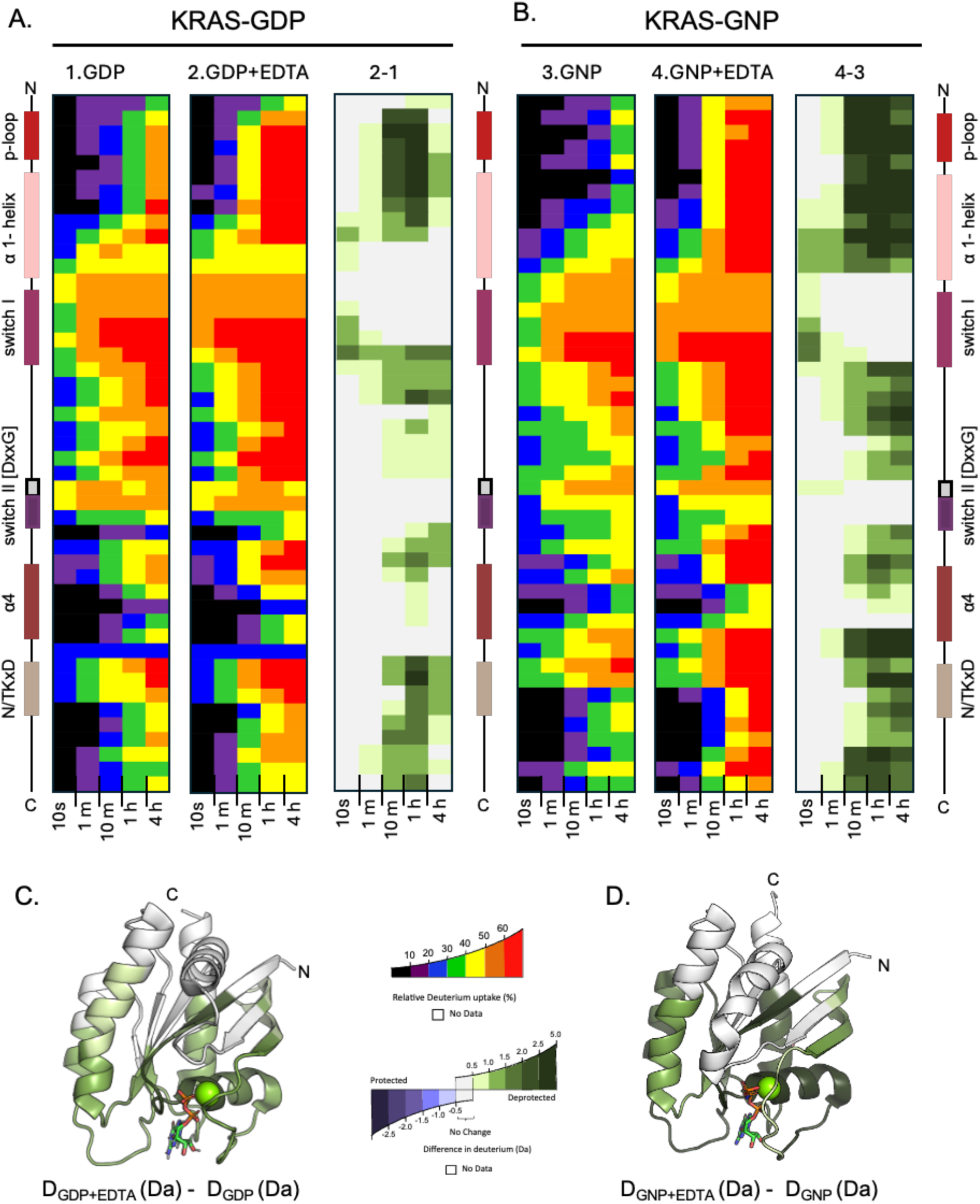
Conformational dynamics of KRAS in different states using HDX-MS. HDX are represented by relative percent deuterium for each peptide in the form of chicklet plots for KRAS in its (A) inactive (GDP-bound) panel 1,2 and (B) active state (GNP-bound) panel 3,4 each in two different conditions, -/+ EDTA treated. The deuterium difference in Da is calculated by subtracting EDTA treated KRAS from untreated KRAS. Two-dimensional representation of KRAS is given in a linear fashion from N terminus (top) to C terminus (bottom), and the locations of key structural elements are shown on the left. All the deuterium-labeling time points are shown, increasing from left to right. All the color scales are shown below. (C and D) Cartoon representation of KRAS-GDP, PDB: 4OBE (C) and KRAS-GNP, PDB: 6OB2 (D) with HDX-MS differences at labeling time point 10 min (upper panel) annotated using the color scheme shown below.

The visual inspection of the differential deuterium uptake calculated between Mg^2+^-bound and Mg^2+^-free states (Figure 1A, panel 3 and Figure 1B, panel 3) revealed widespread increases in deuterium uptake upon Mg^2+^ depletion, indicating enhanced backbone dynamics across multiple KRAS regions. The most pronounced differences appeared in the p-loop, α1-helix, switch I, α4-helix, nucleotide-binding region, and C-terminal region; notably, switch II showed no significant change. Overlapping peptides from the p-loop and α1-helix exhibited an average increase in deuterium incorporation of +5.2 Da relative to Mg^2+^-bound KRAS (Supplemental Figure S1A). Strikingly, these regions displayed evidence of non-EX2 HDX kinetics—a hallmark of slow, correlated conformational transitions manifested as bimodal isotopic envelopes in the raw mass spectra. Each envelope corresponds to a distinct structural population: in this case a protected “closed” state (exchange-incompetent; blue shading in Figure S2A) and an exposed “open” state (exchange-competent; green shading in Figure S2A) [32]. Mg^2+^-bound KRAS–GDP and KRAS–GMPPNP exhibited predominantly closed populations with only approximately 15% of open-state occupancy at the 10-minute labeling point. Whereas Mg^2+^-depleted KRAS showed a dramatic shift toward the open population in both the p-loop and α1-helix—approximately 75% of open-state occupancy at the same time point (Supplemental Figure S2A). This transition demonstrates the profound destabilizing effect of Mg^2+^ depletion, driving KRAS into a markedly more dynamic conformation.

The structural basis for these observations is straightforward: the p-loop and α1-helix lie adjacent to the Mg^2+^-binding site, with Ser17 of the p-loop directly coordinating the metal ion, while switch I residues Tyr32, Asp33, Pro34, and Ile36 contribute to the coordination environment [23]. Consequently, Mg^2+^ depletion directly perturbs both the p-loop and switch I, disrupting local structure and increasing backbone flexibility. The concomitant HDX increases at distal sites—the α4-helix, nucleotide-binding region, and C-terminus—likely reflect long-range allosteric coupling within the KRAS architecture. Importantly, similar differential HDX signatures appeared for KRAS in both GDP- and GMPPNP-bound states upon Mg^2+^ removal, revealing that Mg^2+^ plays a critical role in preserving structural integrity regardless of nucleotide state. Collectively, these results establish Mg^2+^ as a key stabilizing cofactor essential for maintaining both inactive and active KRAS conformations.

To determine whether this Mg^2+^-dependent stabilization is preserved in clinically relevant mutants, we performed analogous HDX-MS experiments on the KRAS G12D variant (Supplemental Figure S3). Comparison of Mg^2+^-bound KRAS G12D-GDP to wild-type KRAS-GDP revealed decreased deuterium incorporation in the N-terminal region (p-loop and α1-helix). More importantly, comparing Mg^2+^-bound versus Mg^2+^-depleted KRAS G12D-GDP revealed a pattern of Mg^2+^-dependent conformational changes similar to wild-type protein. This result revealed that, despite its oncogenic mutation, KRAS G12D retains fundamental reliance on Mg^2+^ for maintaining its native structural integrity and stability of the nucleotide-binding pocket.

### Mg^2+^ Concentration Dependence of KRAS Conformational Dynamics

To investigate how Mg^2+^ concentration modulates the KRAS native conformational state, we performed HDX-MS titration experiments using our Mg^2+^-depleted KRAS. We selected a 10-minute pulse labeling time based on the non-EX2 like bimodal isotopic distributions observed in the p-loop and α1-helix, which exhibited high sensitivity to Mg^2+^ depletion (Figure 3A). Mg^2+^ concentrations ranging from 0.25 μM to 2.5 mM were titrated against 30 μM Mg^2+^-free KRAS, and the resulting HDX profiles were compared to fully native Mg^2+^-bound KRAS.

**Figure 3:**
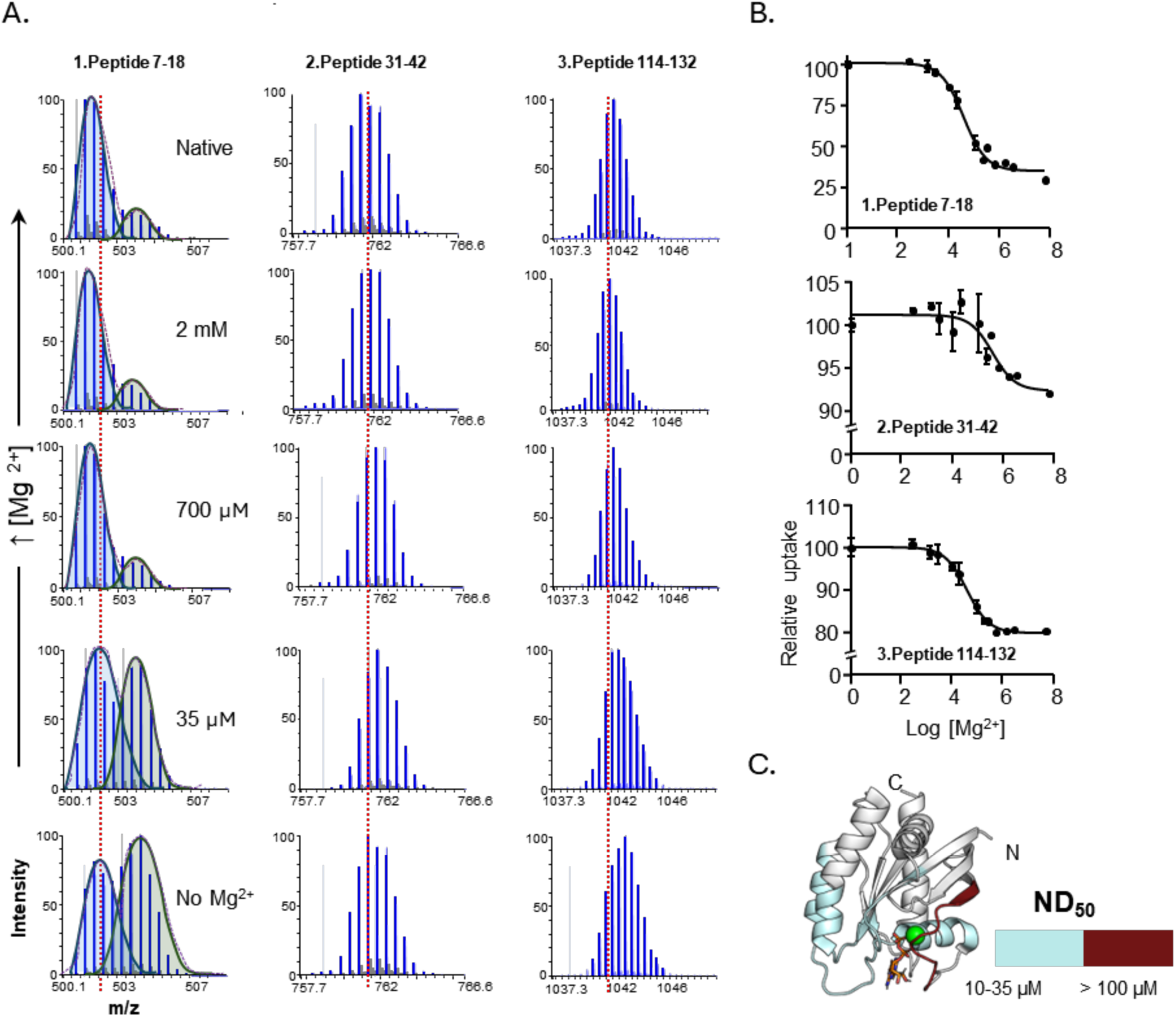
KRAS structural dynamics monitored by HDX-MS by titrating Mg^2+^ at different concentrations. (A) Examples of peptide specific HDX-MS data with representative mass spectra of EDTA treated KRAS show the gradual decrease in deuterium uptake with increase in [Mg^2+^] at 10 min time point and is compared with the topmost mass spectra of no EDTA treated KRAS (Native). Panel 1, peptide 7-18 a representative peptide from the p-loop showing non-EX2 like kinetics with two populations colored blue and green. Panel 2, peptide 31-42 representative peptide from switch I. Panel 3, peptide 114-132 representative peptide from nucleotide binding region. Different [Mg^2+^] shown increasing from bottom to top on the right. The x-axis represents the m/z ratio and the y-axis shows the intensity. Centroid of the mass spectral envelope is indicated with dashed lines (red). (B) A concentration response curve for EDTA-treated KRAS (30 µM) for representative peptides in the presence of different [Mg^2+^]. X- axis shows the log of [Mg^2+^] and y-axis represents the relative percent deuterium uptake. (C) Cartoon representation of KRAS-GDP, PDB ID: 4OBE, with Native dynamics 50% (ND 50) mapped onto the structure according to the color scheme shown below. (C and D) Cartoon representation of KRAS-GDP, PDB: 4OBE (C) and KRAS-GNP, PDB: 6OB2 (D) with HDX-MS differences at labeling time point 10 min (upper panel) annotated using the color scheme shown below.

Quantitative analysis revealed progressive reduction in deuterium uptake across multiple structural regions—p-loop, α1-helix, switch I, α4-helix, nucleotide-binding region, and C-terminal region—as Mg^2+^ concentration increased (Figure 3A). For each structural element shown in Figure 1, at least two representative overlapping peptides were used to calculate the Mg^2+^ concentration corresponding to 50% restoration of native structural dynamics (ND_50_). Data were fit to a non-linear regression model using GraphPad Prism using relative deuterium values; derived ND_50_ values are summarized in Table 1.

**Table 1:** Measuring [Mg^2+^] dependence of KRAS structural elements.

| Structural Elements | Overlapping Peptides | No of a.a residue | ND 50% (μM) | Avg (μM) | SD |
| --- | --- | --- | --- | --- | --- |
| p-loop | 7-18 | 12 | 31.53 | 30.15 | ± 1.84 |
|  | 7-16 | 10 | 28.77 |  |  |
| α1- helix | 17-25 | 9 | 23.46 | 22.48 | ± 1.13 |
|  | 17-23 | 7 | 21.51 |  |  |
| SW-I | 31-42 | 12 | 295.01 | - | - |
| α2-helix | 82-92 | 11 | 25.06 | 27.82 | ± 4.08 |
|  | 83-92 | 10 | 30.59 |  |  |
| NBR | 114-132 | 19 | 34.84 | 31.91 | ± 4.20 |
|  | 114-136 | 23 | 28.98 |  |  |

Surprisingly, four of the five structural elements—all except switch I—regained near 50% of native backbone dynamics at 22-32 μM Mg^2+^ (Figure 3B, Figure S3) and approached complete restoration at 175 μM. This effect was particularly evident in the p-loop, which exhibited non-EX2 kinetics visible in raw mass spectra as a distinct open, exchange-competent population (green shading with higher m/z values; Figure 3A, residues 7–18). As the concentration of Mg^2+^ increased, the amount of this population diminished while the closed, exchange-incompetent population grew, achieving nearly native dynamics at 700 μM Mg^2+^. In striking contrast, switch I exhibited the highest ND_50_ (295 μM) and required 1.25 mM Mg^2+^ to fully regain native conformation (Figure 3A, residues 31–42; Figure 3B), indicating exceptional sensitivity to Mg^2+^ availability. At 1.25 mM Mg^2+^, the overall deuterium uptake pattern of switch I closely matched the native Mg^2+^-bound state, suggesting near-complete recovery of native backbone dynamics.

These results demonstrate that the KRAS conformational equilibrium responds sensitively to free Mg^2+^ concentration, with each structural element displaying a characteristic Mg^2+^ dependence for maintenance of native dynamics. While most structural elements regain native backbone dynamics within the micromolar Mg^2+^ range, switch I requires millimolar concentrations for full recovery. Given that physiological free Mg^2+^ levels typically range from 0.8–1.2 mM [6], our data imply that complete native dynamics are achieved only when Mg^2+^ concentrations approach the upper end of this physiological window. The requirement for elevated Mg^2+^ to restore switch I dynamics reflects a finely tuned balance between backbone flexibility and structural integrity—enabling switch I to fulfill its essential role as a regulatory hub recognized by multiple downstream effector proteins [31].

### SOS_cat_-Mediated Activation of KRAS Involves Mg^2+^-Sensitive Conformational Dynamics

Because cellular free Mg^2+^ concentrations are maintained at approximately 0.8–1.2 mM, complete Mg^2+^ depletion sufficient to drive nucleotide exchange is unlikely to occur *in vivo*. Instead, the cellular machinery responsible for promoting exchange is SOS1[24]. Structural characterization and site-directed mutagenesis studies have established that the SOS1 catalytic domain (SOS_cat_) directly contacts KRAS-GDP through switch I and switch II [6] [24]. To understand how KRAS structural dynamics change upon SOS1 engagement, we investigated conformational changes induced by SOS_cat_ binding using HDX-MS, comparing recombinant KRAS–GDP in solution with and without excess SOS_cat_.

Comparison of HDX profiles revealed localized differential exchange throughout KRAS upon SOS_cat_ binding. Importantly, increased HDX appeared in the same structural elements affected by Mg^2+^ depletion: the p-loop, nucleotide-binding region, α1-helix, and C-terminal region (Figure 4A and Figure 1A). In contrast, switch I exhibited *decreased* deuterium incorporation in the presence of SOS_cat_ (Figure 4A, panel 1). The regions showing increased HDX likely reflect enhanced backbone flexibility upon SOS_cat_ engagement—dynamics that may be essential for GDP displacement. Conversely, reduced HDX in switch I is consistent with direct stabilization from SOS1 [24], which restricts local motion even as other regions become more dynamic.

**Figure 4:**
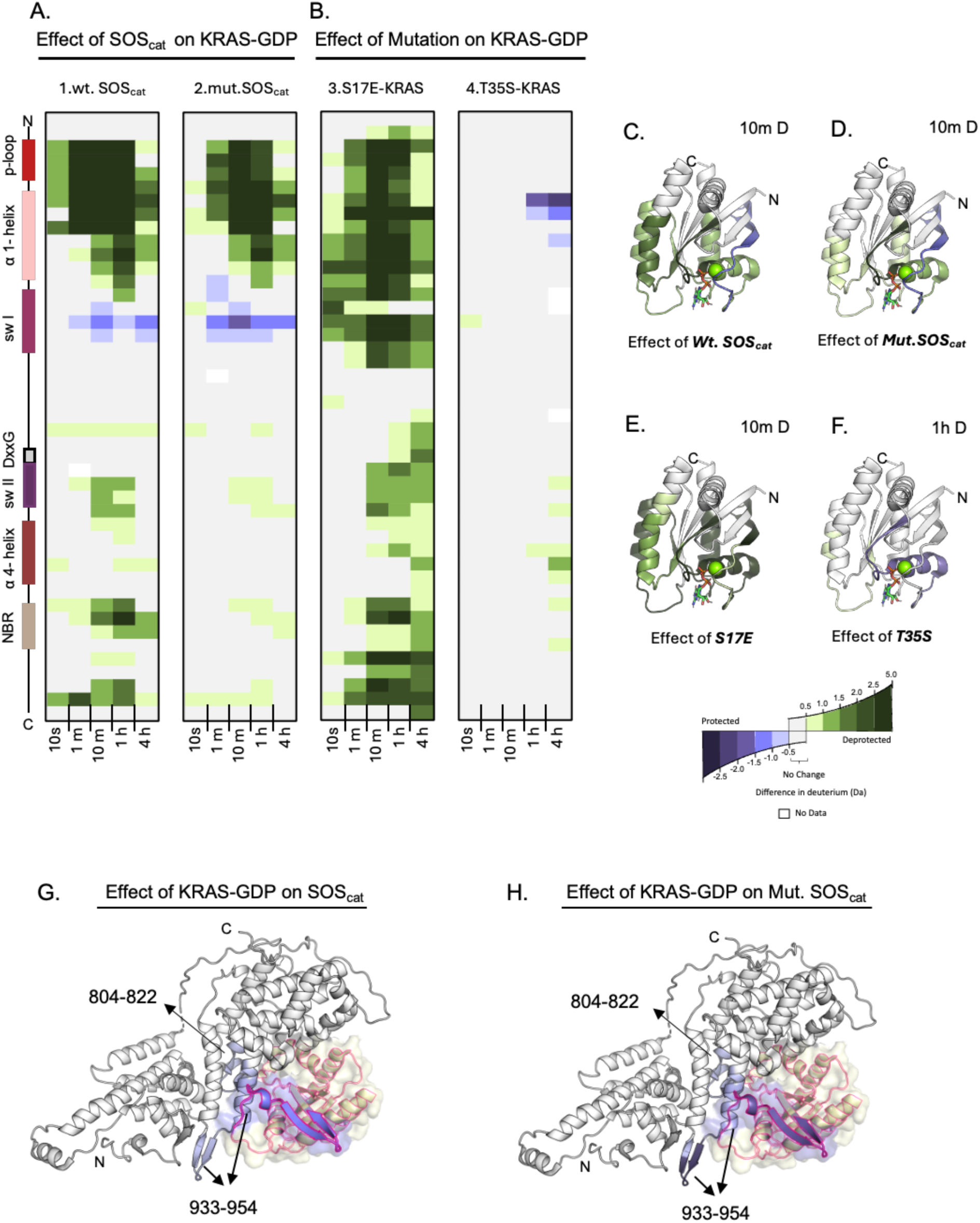
Conformational response of KRAS upon its interaction with GEFs or impact of single point mutation. Differences in HDX are represented by relative deuterium levels of WT KRAS-GDP (A, 1) in the presence of excess WT-SOS_cat_ minus WT KRAS-GDP alone (A, 2) in the presence excess of L938A/E942A SOS_cat_ (Mut. SOS_cat_) minus WT KRAS-GDP alone (B, 3) S17E KRAS variant minus WT KRAS-GDP (B, 4) T35S KRAS variant minus WT KRAS-GDP, color scale shown at the bottom. Two-dimensional representations of WT KRAS are given in linear fashion from N terminus (top) to C terminus (bottom), and the locations of key structural elements are shown on the left. All deuterium-labeling time points are shown, increasing from left to right. (C, D, E and F) Cartoon representations of KRAS-GDP, PDB: 4OBE, with HDX-MS differences at labeling time point 10 min (C) panel 1 (D) panel 2 (E) panel 3 and (F) panel 4 at labeling time point 1 h (upper panels) annotated using the color scheme shown below. (G and H) Cartoon representation of SOS_cat_ bound to KRAS, PDB ID: 7KFZ, with HDX-MS differences at labeling time point 5 s. Differences in HDX are represented by relative deuterium levels of SOS_cat_ (H) WT SOS_cat_ in the presence of KRAS minus WT SOS_cat_ alone (I) L938A/E942A SOS_cat_ in the presence of KRAS minus L938A/E942A SOS_cat_ alone. The transparent surface structure of KRAS with the cartoon representation of PDB ID: 7KFZ, is shown with the red outline with HDX-MS differences at labeling time point 10 m for KRAS from upper panel.

AS our HDX-MS analyses suggested that Mg^2+^ coordination disruption is essential for SOS1-mediated structural changes enabling nucleotide exchange, we examined binding of a previously characterized SOS_cat_ double mutant (L938A/E942A) to KRAS [33]. X-ray crystallography identified Leu938 and Glu942 as key residues that directly contact both Mg^2+^ and the phosphate group of bound nucleotide [34]; mutating these residues impairs the ability of SOS_cat_ to disrupt Mg^2+^ coordination [33].

HDX-MS analysis of KRAS in the presence of excess L938A/E942A SOS_cat_ (Figure 4B, panel 2) revealed increased deuterium incorporation similar to wild-type SOS_cat_, but with considerably smaller magnitude. Interestingly, the mutant produced a larger decrease in HDX at switch I compared to wild-type protein, suggesting a more stable binding interaction with reduced overall conformational perturbation. These findings indicate that while limited conformational rearrangement can occur upon L938A/E942A SOS_cat_ interaction, effective Mg^2+^ perturbation by SOS1 might be required to achieve the full conformational transition necessary for efficient nucleotide exchange.

We also performed HDX-MS on SOSand the L938A/E942A mutant in the presence and absence of excess KRAS to monitor SOS_cat_ structural dynamics upon KRAS binding (Figure 4G, H). In both constructs, decreased deuterium uptake appeared in regions spanning residues 804–822, 933–954, and 960–981(Supplemental datafile, tab Fig4). These regions comprise part of the helical hairpin motif within the CDC25 domain of SOS1, previously identified through structural and molecular dynamics studies as the interface engaging switch I during nucleotide exchange [35]. The observed HDX differences at these regions demonstrate direct KRAS binding and support their functional role in SOS1-mediated nucleotide exchange.

### Disruption of Mg^2+^ Coordination Differentially Modulates KRAS Conformational Dynamics

To further probe the importance of Mg^2+^ in maintaining KRAS conformational dynamics, we examined two separate point mutants: the phosphomimetic KRAS S17E and the mechanistic variant KRAS T35S. The S17E substitution was designed to perturb the Mg^2+^ coordination environment: because Mg^2+^ forms a coordination bond with Ser17 in the p-loop, substituting glutamate should disrupt Mg^2+^ coordination geometry and promote alterations in backbone dynamics similar to those observed upon Mg^2+^ depletion or SOS_cat_ interaction.

HDX-MS analysis of the S17E KRAS variant compared to Mg^2+^-bound wild-type KRAS confirmed these predictions (Figure 4B panel 3). The S17E KRAS variant exhibited increased deuterium incorporation throughout the protein backbone, with pronounced effects in the p-loop, α1-helix, switch I, nucleotide-binding region, α4-helix, and C-terminal region—changes exceeding even those observed with EDTA-treated KRAS (Figure 2A). These findings support a model wherein perturbation of Mg^2+^-coordinating interaction shifts KRAS away from its native state, highlighting the critical role of Mg^2+^ coordination integrity in maintaining native KRAS architecture.

Next, HDX MS analysis T35S mutant provided complementary insights. While Thr35 directly coordinates Mg^2+^ in the active GTP-bound state to stabilize switch I in a closed, signaling-competent conformation, its role in the inactive GDP-bound state is less defined. Substitution of Thr35 with serine disrupts Mg^2+^ coordination in KRAS-GTP, forcing a “State 1” conformation that mimics the GDP-bound state and produces partial loss of function [36, 37]. Our investigation focused on the GDP-bound state. Interestingly, the T35S mutation resulted in *decreased* HDX compared to wild-type KRAS-GDP in the p-loop and α1-helix (residues 7–30), alongside subtle HDX increases in the nucleotide-binding region at later labeling time points (Figure 4B, panel 4). These observations suggest that although Thr35 does not reside within the primary Mg^2+^ coordination sphere in the GDP-bound state—where two water molecules typically occupy the Thr35 position—it nonetheless influences the structural dynamics of the surrounding environment.

### Functional Evidence Linking Mg^2+^ Perturbation to SOS1-Mediated KRAS Activation

To functionally validate our structural findings, we performed nucleotide exchange assays comparing GDP-to-GTP exchange rates for wild-type KRAS and the S17E KRAS variant. For wild-type KRAS, assays were conducted independently with the wild-type and the L938A/E942A mutant of SOS_cat_; the S17E KRAS variant was assayed with wild-type SOS_cat_. The highest nucleotide exchange rate occurred with wild-type KRAS and wild-type SOS_cat_, consistent with the GEF’s ability to perturb Mg^2+^ coordination and promote nucleotide release (Figure 5A). In contrast, the L938A/E942A SOS_cat_ mutant—deficient in disrupting Mg^2+^ coordination—failed to promote nucleotide exchange (Supplemental Figure S4) emphasizing the exquisite structural requirements of this protein-protein interaction. Consistent with our HDX-MS data showing widespread increases in dynamics, the S17E KRAS variant displayed a significantly elevated intrinsic nucleotide exchange rate relative to wild-type KRAS; this rate was further accelerated upon addition of wild-type SOS_cat_ (Figure 5A). Collectively, these functional data align with our structural findings, demonstrating that proper Mg^2+^ coordination and its role in maintaining structural integrity are primary determinants of efficient nucleotide cycling.

**Figure 5:**
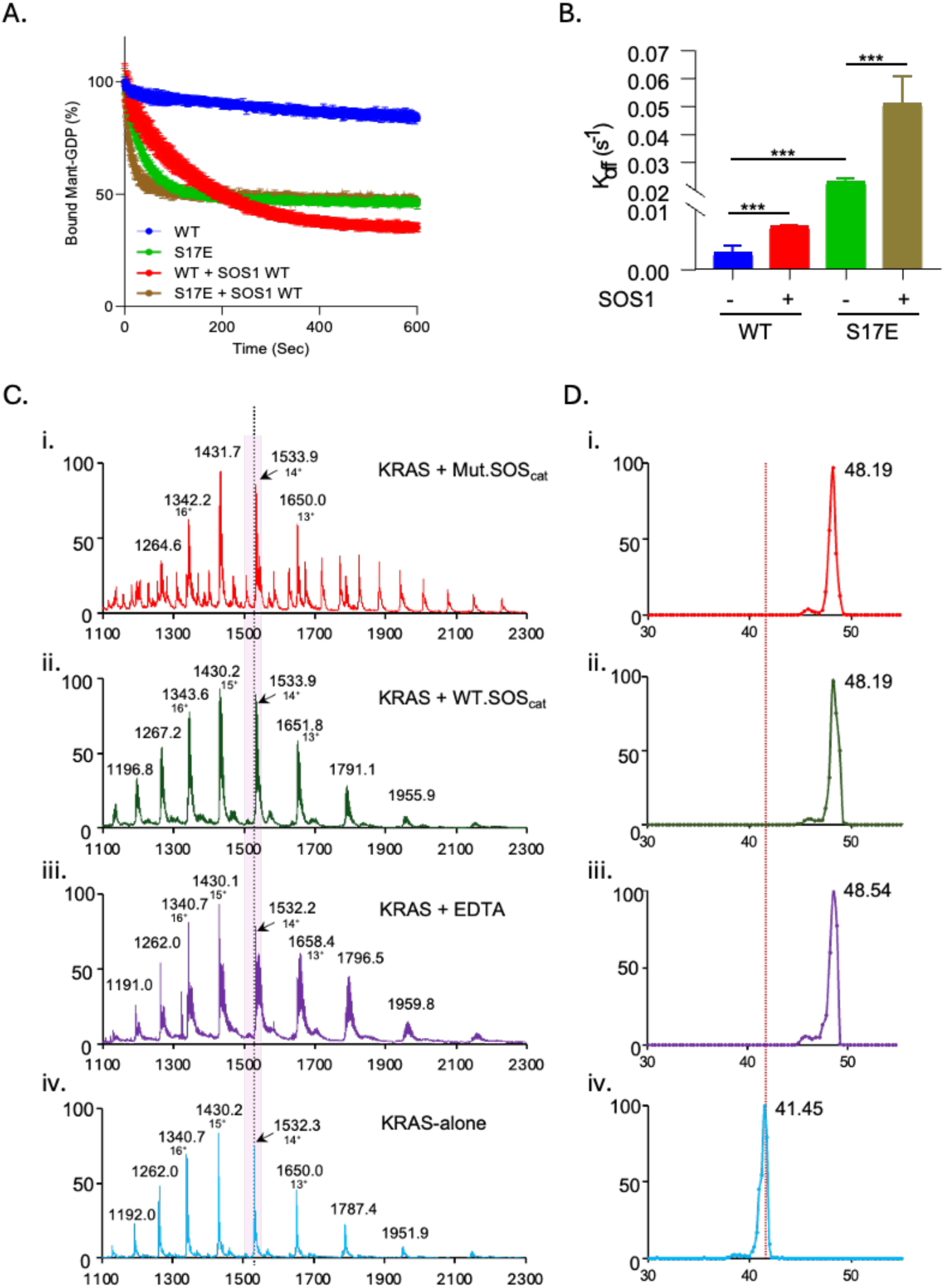
(A and B) Nucleotide exchange assay. Fluorescence-based measurements were used to determine nucleotide exchange kinetics. (A) Time-resolved GDP dissociation from WT KRAS and the S17E mutant in the absence or presence of SOS_cat._ (B) Quantification of nucleotide exchange rates. The intrinsic exchange rate of KRAS S17E is significantly higher than KRAS WT, and it is further enhanced by SOS1. ***, P < 0.001, unpaired t test. (C and D) Global KRAS conformation at different conditions monitored using IM-MS. (C) Raw mass spectra of KRAS (i) in the presence of L938A/E942A SOS(ii) in the presence of WT SOS(iii) KRAS treated with EDTA (iv) untreated KRAS (Native). X-axis shows the m/z ratio, and the y-axis represents the intensity. The pink shaded region highlights the peak corresponding to the 14^+^ charge state. The dashed line indicates the centroid position of that charge state. (D) Ion mobility arrival time distributions for the 14^+^ charge state for respective states which are color coded matching the mass spectra. X-axis shows the drift time in ms and the y-axis represents the intensity.

### Native Ion Mobility Mass Spectrometry Reveals Global Conformational Changes Upon Mg^2+^ Depletion and SOS1 Interaction

Complementing our HDX-MS analysis, we employed native ion mobility mass spectrometry (native-IMS) to assess whether the observed local changes in backbone dynamics correspond to global conformational change. While our earlier native-MS experiments (Figure 1) provided accurate mass measurements of KRAS–nucleotide complexes using time-of-flight detection, ion mobility adds a second dimension of separation: molecular ions of identical m/z are separated based on drift time, with compact, globular conformations traveling faster than extended or open structures [38].

Native-IMS experiments were conducted on a Waters SELECT Series Cyclic IMS platform under near-native conditions (50 mM ammonium acetate, pH 7.8). Consistent with HDX-MS results, EDTA-treated KRAS, SOS_cat_-bound KRAS, and L938A/E942A SOS_cat_-bound KRAS all exhibited increased drift times (48.54 ms) relative to Mg^2+^-bound wild-type KRAS (41.54 ms) (Figure 5D). This shift indicates a transition from a compact shape—observed in Mg^2+^-bound KRAS—toward a more open, extended conformation upon EDTA treatment or SOS_cat_ binding (Figure 5D). The similarity in drift times among Mg^2+^-depleted and SOS_cat_-bound states reinforces that both conditions induce similar levels of global structural expansion, albeit with varying magnitudes of local flexibility as observed by HDX-MS. These results demonstrate that Mg^2+^ coordination plays a critical role in maintaining the global architecture of native KRAS, and its loss or perturbation shifts the conformational ensemble toward a more dynamic, open state.

## Discussion

Previous structural and computational studies—encompassing molecular dynamics simulations, X-ray crystallography, and NMR relaxation analyses—have highlighted the critical role of Mg^2+^ in stabilizing the KRAS–nucleotide complex, enhancing nucleotide affinity, and enabling SOS1-mediated exchange [22, 23, 31, 39–44]. The present study extends these findings by delineating the structural and mechanistic contributions of Mg^2+^ to stabilizing the native conformational ensemble of KRAS and facilitating its transition between inactive GDP-bound and active GTP-bound states. Although Mg^2+^ is known to coordinate the phosphate moiety of bound nucleotide and hydroxyl groups of serine and threonine residues, our results demonstrate that its role extends far beyond simple coordination. We show that Mg^2+^ functions as a conformational gatekeeper: its removal compromises the structural integrity of key regulatory elements including the p-loop, α1-helix, switch I, nucleotide-binding region, α4-helix, and C-terminal region (Figure 1). Perturbation of Mg^2+^ coordination whether through chelation, Ser17 mutation, or SOS_cat_ engagement, shifts the protein toward a more open, exchange-competent ensemble.

Switch I and switch II form the binding interface for downstream effector proteins in KRAS–GTP [31]. As switch I serves as a primary interface for effector recognition, its marked dependence on millimolar Mg^2+^ levels underscores the importance of Mg^2+^ in preserving signaling accuracy. Our findings reveal that the KRAS nucleotide cycle depends not only on GEF and GAP activities but also on the Mg^2+^-dependent stability of its structural framework.

In the cellular context—where intracellular Mg^2+^ concentrations (approximately 0.8–1.2 mM) are substantial and GTP levels exceed GDP by roughly tenfold—complete Mg^2+^ removal by SOS1 during catalysis is unlikely [45]. Instead, the SOS1 catalytic domain likely induces a transient distortion in Mg^2+^ coordination sufficient to lower nucleotide affinity and permit GDP dissociation with GTP insertion from the cellular pool. Our Mg^2+^ titration data establish switch I as the most Mg^2+^-dependent region of KRAS, requiring the upper end of the physiological Mg^2+^ range to fully regain native structural dynamics. Importantly, this same region exhibits decreased deuterium incorporation upon SOS_cat_ binding (Figure 4). This dual behavior—high Mg^2+^ sensitivity coupled with protection from exchange upon SOS1 engagement—supports a model wherein SOS1-mediated opening of KRAS for nucleotide exchange must be precisely tuned. Prior RAS–SOS1 structures show that KRAS at the SOS1 catalytic site is devoid of Mg^2+^, supporting the notion that SOS1 perturbs Mg^2+^ coordination as part of the exchange mechanism [24, 46]. By perturbing Mg^2+^ coordination sufficiently to weaken GDP binding while simultaneously stabilizing switch I, SOS1 prevents structural collapse during the transient nucleotide-free intermediate. Thus, SOS1 functions not only as a catalytic GEF but also as a molecular chaperone, holding switch I in a productive conformation while generating the Mg^2+^-perturbed, exchange-competent state required for efficient KRAS activation.

Furthermore, KRAS S17E mutation has been described as a sensitizing variant [47], a phenotype readily rationalized by its impaired Mg^2+^ coordination. Because Ser17 directly contributes to Mg^2+^ binding, glutamate substitution disrupts the octahedral geometry required for stabilizing nucleotide-bound states. Our HDX analysis reveals that S17E exhibits widespread increases in backbone dynamics compared to wild-type KRAS and undergoes intrinsic GDP-to-GTP exchange. By perturbing Mg^2+^ coordination at Ser17, the S17E variant fails to stabilize either nucleotide-bound state, instead sampling a hyperdynamic, partially unstructured conformational ensemble. This instability likely prevents proper adoption of the GTP-bound active conformation required for effector recognition and downstream signaling. Despite increased mobility, S17E KRAS is functionally compromised—leading to enhanced susceptibility due to impaired signaling competence, consistent with its classification as a sensitizing mutant [47]. These observations highlight the delicate balance between flexibility and structural stability that depends on Mg^2+^ coordination: while limited perturbation, as mediated by SOS1, is necessary for exchange, complete disruption as in S17E abolishes the ability to achieve a stable, signaling-competent state. Building on these mechanistic insights, future drug-discovery efforts may exploit the dynamic allosteric landscape of KRAS revealed here (Figure 6). Systematic fragment-based and computational screening targeting transiently exposed pockets—particularly around the Mg^2+^ coordination network—could identify ligands that either reinforce or disrupt local structural stability. The dynamic regions defining these allosteric nodes might serve as anchor points for PROTAC-style degraders or conformation-selective stabilizers, offering strategies to trap KRAS in an inactive GDP-like state or destabilize the active conformation altogether. Such approaches emphasize that targeting conformational plasticity—not just catalytic pockets—represents a promising avenue for next-generation KRAS therapeutics.

**Figure 6:**
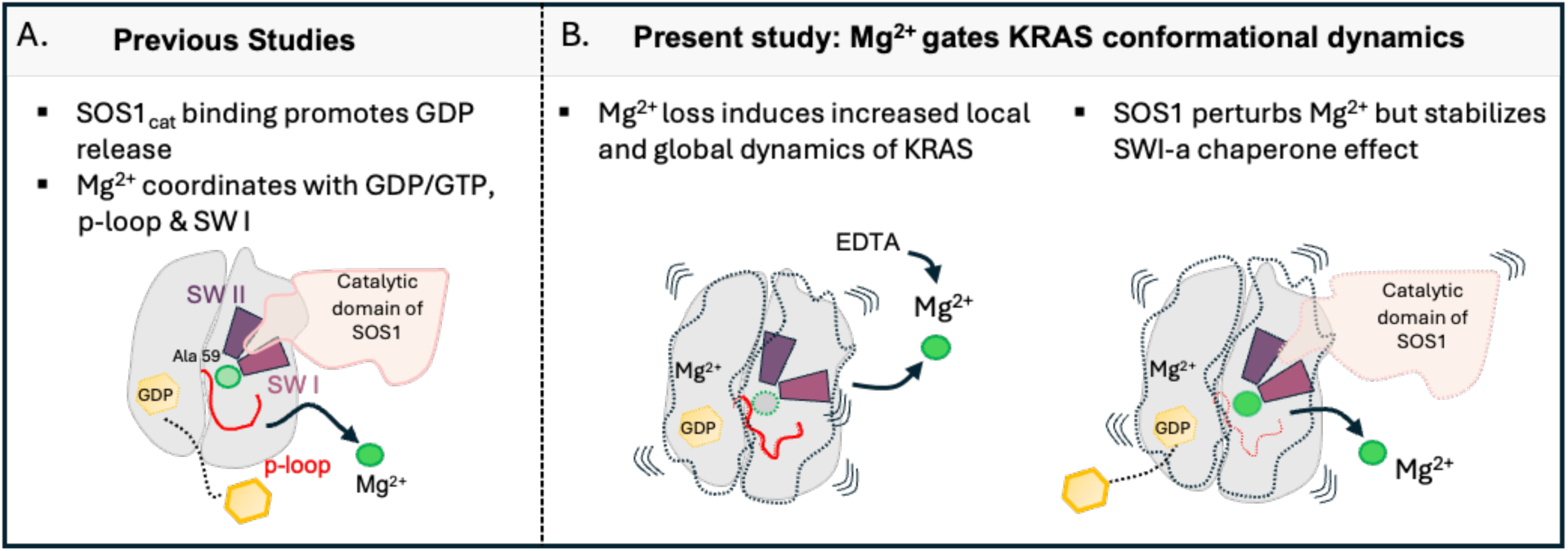
Mg^2+^ functions as a conformational gatekeeper of KRAS. The schematic illustrates: (A) Previous Study: Earlier research established that SOS1 promotes nucleotide exchange by engaging switches I and II and also emphasizing the importance of localized Mg^2+^ coordination at the active site. Studies primarily are static snapshots. (B) Current Findings: Using HDX-MS, we demonstrate that disrupting Mg^2+^ coordination, whether through EDTA chelation, SOS1 binding, or targeted mutation, induces widespread, long-range conformational dynamics that extend far beyond the Mg^2+^-binding site. These Mg^2+^-dependent structural dynamics of KRAS reveal that Mg^2+^ is far more than a simple cofactor; it functions as a central regulator of protein-wide stability.

## Methods

### Protein expression, purification, and nucleotide exchange

The gene encoding human WT-KRAS (residues 1-169) or WT-SOS_cat_ (558-1049, Uniprot Q07889) was synthesized and inserted into a plasmid to introduce an N-terminal His tag for affinity purification (Genescript). Constructs clonesd in pET 28a(+) were expressed using *E. coli* BL21 (DE3) cells. Starter cultures were grown overnight in LB media containing 0.1 mg/mL kanamycin. Overnight cultures were diluted 50-fold and grown at 37°C to A_600_ 0.5, then induced with 0.25 mM IPTG for overexpression of protein at 30°C for 4 h with shaking. The cells were then pelleted using centrifugation at 7000 RPM for 15 mins. The collected pellet was then resuspended in lysis buffer containing 20mM Tris HCl, pH 7.6, 250 mM NaCl, 20 mM imidazole and 2.5 mM PMSF and then subjected to sonic dismembranation ice. The lysate was then clarified using high speed centrifugation at 19000 RPM for 1 h. The resultant supernatant was used for further purification using Nickel affinity chromatography using standard protocols. The protein was eluted with 250 mM imidazole containing buffer. The eluted protein was then subjected to size-exclusion chromatography using Superdex 75 10/300 GL column (GE Healthcare Life sciences). Fractions containing pure protein were collected, concentrated, and the concentration was confirmed using Bradford assay.

In order to switch nucleotides for both native MS and HDX MS analyses, 200 µL of WT-KRAS was treated with an excess concentration of 2.5 mM EDTA to chelate the bound Mg^2+^ and incubated with gentle rotation at room temperature for 45 min to ensure complete Mg^2+^ removal. Following incubation, an excess of GMPPNP was added to the reaction mixture, and the sample was placed on ice for an additional 30 min to facilitate nucleotide exchange. The sample was then divided into two equal aliquots of 100 µL each for differential Mg^2+^ treatment. Using spin desalting columns, one aliquot was buffer-exchanged into a no-magnesium buffer (150mM NaCl, 20mM HEPES), while the second aliquot received 2.8 mM MgCl₂, was incubated for 10 min to allow Mg^2+^ re-binding to the KRAS–GMPPNP complex and was subsequently buffer-exchanged under the same conditions. All buffer exchanges were performed using identical column (2mL Zeba spin columns, Cat#: 89890, ThermoFisher Scientific) and centrifugation parameters (3,000 rpm for 2 min).

### Native MS

Protein samples were first exchanged into a buffer containing 50 mM ammonium acetate pH 7.8 using 2 mL Zeba (Cat#: 89890, ThermoFisher Scientific) desalting cartridges. Mass spectra of the buffer exchanged samples were acquired using micro-ESI fluidics at 10 µL/min (Waters Synapt G2Si MS or Waters Select Series Cyclic IMS). Mass calibration was performed using major mix (Waters). Spectra were acquired in the m/z range 500-8000 Da in triplicates with following instrument parameters: Capillary voltage 1.4 kV; sampling cone voltage 60 V; source offset 80 V; source temperature 37 °C, Quad profile was set to auto profile. Data was processed with MassLynx software version 4.2. For ion mobility separation of different conformers, IMS wave height was fixed to 40 V and wave velocity was optimized at 900m/s to achieve the best separation of molecules in the mobility cell to record the drift time.

### Nucleotide Exchange assay

KRAS WT and S17E mutant (100 μM) were incubated with 300 μM Mant-GDP (Cat#: NU-204L, Jena Bioscience) in Tris buffer (20 mM Tris pH7.5, 50 mM NaCl, 5 mM EDTA) at room temperature for 2 hours. Reactions were terminated with 10 mM MgCl_2_, and the Tris buffer was exchanged to assay buffer (20 mM Tris pH7.5, 50 mM NaCl, 2 mM MgCl_2_), to remove extra EDTA and unbound nucleotides by using 2 mL Zeba (Cat#: 89890, ThermoFisher Scientific) desalting cartridges. To evaluate the intrinsic nucleotide exchange, 50 μl of KRAS WT and mutant (2 µM) were added to black 384-well plate (Cat#: 164564) and 50 μl of GTP (2 mM) was injected into wells. To examine the effect of SOS1 proteins on nucleotide exchange, SOS1 WT and mutant (2 µM) with GTP (2 mM) were applied. Fluorescence was measured every 0.5 s for 10 minutes at 360nm/440nm (excitation/emission) in a Synergy Neo reader (BioTek). Data were exported and analyzed using GraphPad Prism (GraphPad Software). All measurements were repeated at least three times.

### HDX-MS

Deuterium labelling experiments were performed for different constructs of recombinant KRAS. The GDP and the GMPPNP states of each protein were labelled independently with deuterium, at identical experimental conditions to allow for direct comparisons between each set of the bound forms. Similarly, to monitor the binding effects of SOS_cat_ constructs on KRAS, SOS_cat_ was incubated with KRAS at 5:1 molar ratio and was incubated for 5 mins at room temperature prior to deuterium labelling.

HDX was initiated by diluting 1 µL of protein (35 µM) in buffer (20 mM HEPES, 150 mM NaCl, pH 7.8), 19-fold into labelling buffer (20 mM HEPES, 150 mM NaCl, pD 7.8, 99% D_2_O) at room temperature (23 °C). The labelling reaction was quenched at 5 time points (10s, 1m, 10m, 1h and 4h) using 20 µL of quench buffer (0.8% FA, 0.8 M GdnHCl, pH 2.3) and immediately subjected to LC-MS analyses. All deuterium-labelling experiments were performed in duplicate. Deuterium incorporation was measured as described previously [48]. Briefly, quenched samples were digested online (15°C) using Nepenthesin II (Affipro) and the resultant peptides were desalted using a 1.7 μm BEH trap and then separated via UPLC with a 1 × 50 mm 1.8 μm HSS T3 separation column. All LC components were housed in a cold chamber, maintained at 0°C. Mass spectra were acquired using a Waters Synapt G2Si HDMS^E^ operated in TOF-IMS mode. Peptide masses were identified from PLGS searches using non-specific cleavage of a custom database containing the sequences of human WT-KRAS (residues 1-169) or WT-SOS_cat_ (558-1049, Uniprot Q07889). The relative amount of deuterium in each peptide was determined using DynamX 3.0.1 software by subtracting the centroid mass of the undeuterated form of each peptide from the deuterated form, at each time point, for each condition. The deuterium uptake values (provided in the Supplemental Datafile) were used to generate all difference maps found in the Figures as well as the Supplemental Datafile. Deuterium levels were not corrected for back exchange and thus reported as relative [48]. All HDX-MS data have been deposited to the ProteomeXchange Consortium via the PRIDE [49] partner repository with the dataset identifier PXD074470.

### Quantitative analysis of magnesium titration followed by HDX-MS

Two overlapping peptides were selected from each of the following regions of the KRAS protein: the P-loop, α1 helix, α4 helix, and the nucleotide binding region. One peptide was chosen for the SW1 region. The selected overlapping peptides were not identical in length which may contribute to variability seen by the standard deviations. The relative percent deuterium uptake for each peptide was plotted on the y-axis against the corresponding Mg^2+^ concentrations on the x-axis. The Data visualization and ND_50_ analysis were done using GraphPad Prism. The Mg^2+^ concentrations were log-transformed within Prism using the equation (1) below.

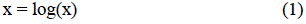

Nonlinear regression fit was applied using the built-in log(agonist) vs. response (three-parameter) model. Separate trend lines were generated for each of the two overlapping peptides per region. ND_50_ was calculated for each peptide and the standard deviations for each region were manually obtained.

## Acknowledgements

Financial support was provided by grant R01 CA-233978 (T.E.W. and J.R.E.) from the National Institutes of Health and CPRIT RP220145 and NIH R01CA244341 (K.D.W).

## Declaration of Interests

K.D.W. has received consulting fees from Sanofi Oncology, Amgen, AstraZeneca, Boehringer Ingelheim and served on the SAB for Vibliome Therapeutics, Vellorum Pharmaceuticals and Low Institute for Therapeutics. K.D.W. has received research funding from Revolution Medicines and Elekta. K.D.W. is co-founder and has equity interest in Stabilix. K.D.W. declares that none of these relationships are directly or indirectly related to the content of this manuscript.

**Supp. Data File 1:** Supplemental figures, Word document containing supplemental figures S1-4.

**Supp. Data File 2:** Excel file containing details of HDX-MS experimental set-up and differences in deuterium incorporation data for all peptides.

## Supplemental Information For

### Contents

Supplementary Figures 1-4

**Figure S1:**
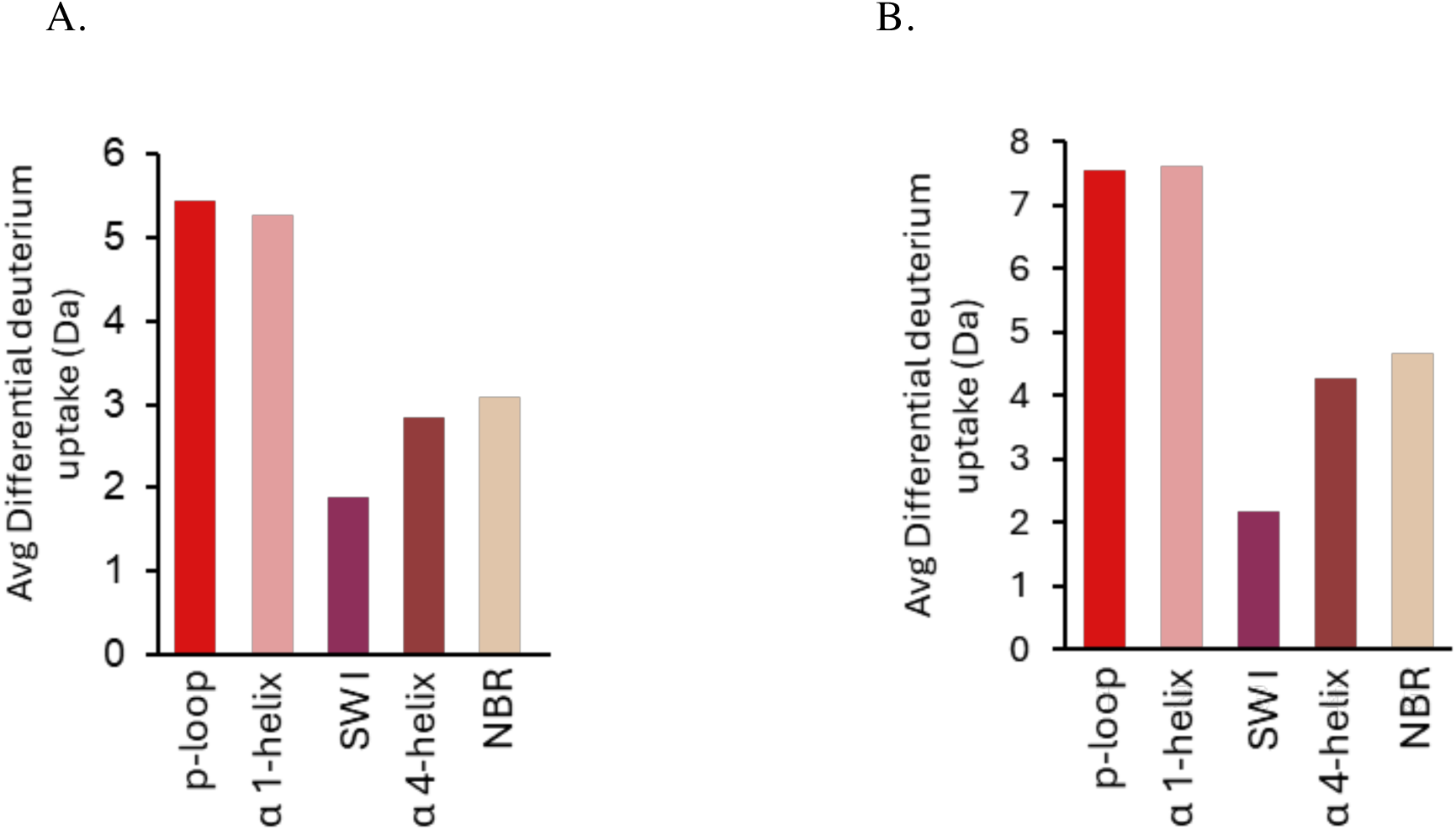
Conformational dynamics of KRAS at key structural elements using HDX MS. The average differential deuterium uptake calculated for each structural elements by taking the overlapping peptides of each region and is shown with different colors in the form of bar graphs. The differential deuterium in Da is calculated (A) KRAS_GDP_-EDTA treated minus KRAS_GDP_ (B) KRAS_GNP_-EDTA treated minus KRAS_GNP_. Y-axis represents the average differential deuterium (Da) and x-axis shows the structural elements.

**Figure S2:**
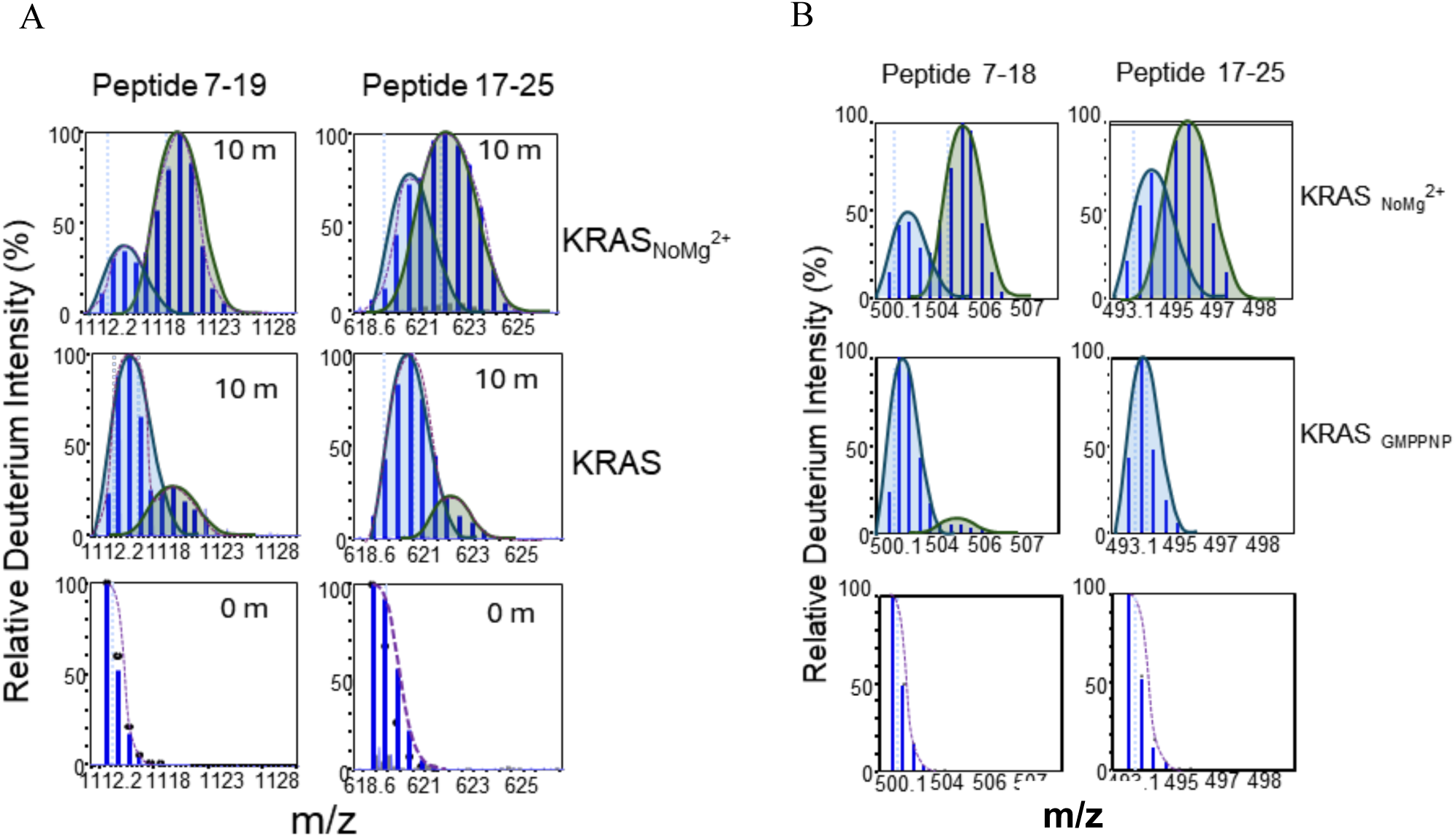
EDTA treated KRAS shows increased dynamics at the p-loop region. Examples of peptide specific HDX MS data with representative mass spectra of EDTA treated KRAS, represented as KRAS_NoMg_^2+^ at the top panel compared to the middle panel which shows the MS spectra of KRAS (Native) at 10 min deuteration time point. Peptide 7-18 a representative peptide from the p-loop and peptide 17-25 from α1-helix region showing non-EX2 like kinetics with two populations colored blue and green for the EDTA treated KRAS. (A) KRAS in GDP bound (Inactive state). (B) KRAS in GNP bound (Active state). Y-axis represents the relative deuterium in percentage and x-axis shows the m/z scale.

**Figure S3:**
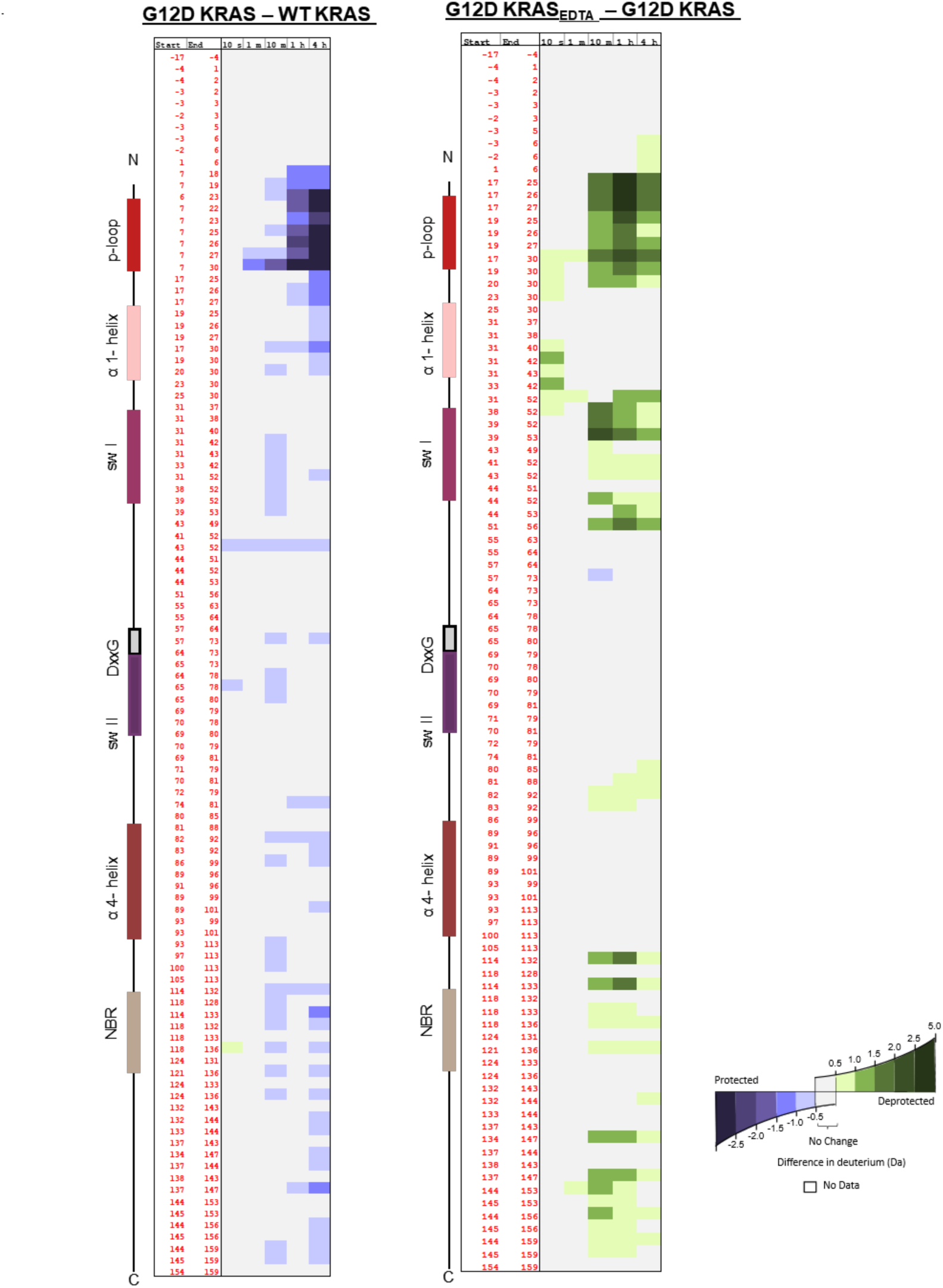
Impact of point mutation at p-loop on the conformational dynamics of KRAS. Differences in HDX are represented by relative deuterium levels of WT KRAS-GDP Panel 1, G12D-KRAS minus WT KRAS-GDP alone. Panel 2 effect Mg^2+^ on G12D KRAS. G12D KRAS_GDP_ EDTA treated minus G12D KRAS_GDP_. Two-dimensional representations of WT KRAS are given in linear fashion from N terminus (top) to C terminus (bottom), and the locations of key structural elements are shown on the left. All deuterium-labeling time points are shown, increasing from left to right. The deuterium difference (Da) is colored according to the color scale shown

**Figure S4:**
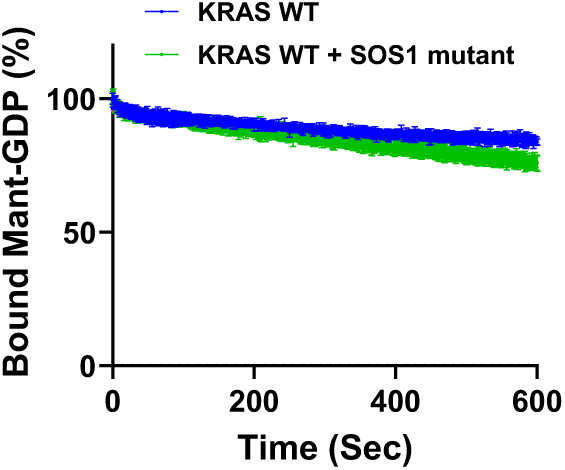
Nucleotide exchange assay. GDP dissociation kinetics from wild-type KRAS were quantified over 10 min with or without the L938A/E942A SOScat variant

## Notes

### Summary of Updates

The manuscript had magnesium written incorrectly; it was shown as a subscript everywhere.

